# Geometry of disordered porous environments regulates cell migration

**DOI:** 10.1101/2025.09.17.676737

**Authors:** Laeschkir Würthner, Frederik Graw

## Abstract

Cell migration is a dynamic process that is of critical importance to various aspects of living organisms, including organogenesis, wound healing, and immune responses. Several external factors are known to influence and direct active cell movement, such as chemokine gradients or the composition and mechanical properties of the extracellular matrix (ECM). While progress has been made in elucidating some of the biochemical pathways that control cell migration, little is known about the impact of the porous structure of the ECM on active cell motion. Here, by combining computational modelling and theory, we reveal how porous environments, as represented by the ECM, determine cell migration dynamics. Simulating cell movement in a 3D cellular Potts model accounting for amoeboid-like cell shape dynamics, we show that cell migration within disordered porous environments is characterized by distinct transient motility regimes that deviate from persistent motion and are best described by ‘hopping’ of cells between ‘traps’. Using theory, we are able to show how these motility regimes and large scale transport properties are linked to geometrical properties of the microstructure. Importantly, our analyses reveal that spatial heterogeneities in the porosity lead to non-homogeneous cell distributions and effectively guide cell movement towards regions of low porosity, an effect which we here term as *porotaxis*. Overall, our work reveals the porosity of the ECM as an important control parameter that shapes cell migration and cellular distribution, and provides a conceptual framework to relate experimentally observed cell motility modes to tissue structures and vice versa. This connection between geometry and cell motility could enhance our understanding of how structural elements shape cell migration and tissue organization in various conditions, such as chronic inflammation, immunity, and cancer.

## I. INTRODUCTION

Cell migration plays an essential role during many biological processes. The development and function of immune responses, wound healing and organ development all represent complex dynamical and highly orchestrated processes that strongly rely on cell motility [1–3]. At the cellular level, migration critically depends on the dynamic interplay between the actomyosin complex and the extracellular matrix (ECM). Intracellular actin polarization, emerging through self-organization, drives the formation of cell protrusions which mechanically couple the cell body to the ECM, enabling cells to move forward by exerting traction forces [4–8]. Cell motility is generally controlled and guided by both intrinsic cues, such as cytoskeletal organization and receptor expression (integrins) along the cell membrane [9], and extrinsic cues, including chemokine gradients as well as the composition and mechanical properties of the ECM [7, 10, 11]. Importantly, intrinsic and extrinsic factors are interconnected and can interact in various ways. For instance, it has been shown that the interplay between environmental cues and cell-induced remodeling of the environment leads to physicochemical footprinting, allowing cells to keep a memory of past paths [12]. In addition, cells can utilize their nucleus as a mechanical gauge to explore their local environment, thereby guiding decision-making and identifying the path of least resistance [13].

Cell motility can be categorized into distinct mechanistic modes, most prominently *mesenchymal* and *amoeboid* migration [7]. Mesenchymal migration is characterized by pronounced protrusions and strong adhesion to the ECM, allowing cells to move forward by actively remodeling their microenvironment through mechanical deformations and proteolysis of the ECM. In contrast, amoeboid migration relies on low adhesion and dynamic cell shape changes that allow cells to squeeze through small constrictions. These mechanistic differences, in combination with environmental constraints and biochemical cues, can manifest in different motility regimes. For instance, it has been shown that cytotoxic CD8^+^ T cells migrate within collagen networks via a two-state persistent random walk, alternating between a slow and fast state [14], whereas other work has shown that their motility in the infected mouse brain follows a Lévy walk (i.e., superdiffusion) [15]. Further examples include lymphocyte migration in lymph nodes, where previous work found location-dependent diffusion regimes, ranging from normal (Brownian) diffusion [16] to anomalous diffusion [17], as well as cancer cell migration in collagen networks, which follows an anisotropic persistent random walk [18].

Taken together, these examples illustrate that cells adopt diverse motility regimes across different tissues, raising the question of what factors ultimately set these modes. While some of the biomolecular pathways that regulate cell migration have been elucidated [19–21], less is known about the role of the cell’s microenvironment in controlling cell movement, in particular the geometric properties of the ECM. The ECM typically exhibits a disordered porous structure, characterized by interconnected heterogeneous pore spaces. The spatial variation in pore size, shape, and connectivity results in complex environments that impose physical constraints on cell motion. Previous experimental studies demonstrated that cell motility is closely linked to these geometric properties [22–25].

Many theoretical approaches have been developed to decipher cell migration in confined environments. Due to the inherent complexity of cell migration and the incomplete understanding of the underlying intracellular processes, theoretical studies frequently employ coarse-grained descriptions, in which cell motion is described as a generalized random walk [26–30]. These coarse-grained models have been shown to reproduce different experimentally observed motilty modes, including superdiffusion, subdiffusion, and persistent random walk. Other studies combined computational models for cell shape dynamics, such as the Cellular Potts model (CPM) and phase-field models, with random walk models to address cell migration within tissues and confined environments [31–35]. More recent studies demonstrated that inference techniques allow to reconstruct the underlying equations of motion for confined cell migration directly from experimental cell trajectories [36–38]. However, since previous approaches typically rely on simplifying assumptions, for example treating cells as spherical objects [28, 39] or considering regular topographic environments [34, 40], it remains unclear how exactly the microenvironment sets the mode of cell motility.

To further elucidate how geometric properties of the ECM influence cell migration, we here investigate the dynamics of motile cells within 3D disordered porous structures. We combine computational modeling with theory to decipher how porous environments control cell motility modes. Specifically, we use a CPM to capture realistic amoeboid-like cell shape dynamics and couple it with active Brownian motion (ABM) [39, 41] to account for cell motility. Simulations of cell movement within the pore space of 3D disordered fibrous porous media, mimicking the natural environment of various cell types, reveal distinct transient motility regimes with transitions from normal to subdiffusive dynamics governed by the porosity of the environment. Our findings imply that cell motility in disordered environments is equivalent to a random walk among traps: cells migrate by hopping from one pore to another and can temporarily become trapped in smaller pores. Using theory, we are able to connect transport properties at large length and time scales to geometric properties of the microstructure. Importantly, our work reveals that spatial heterogeneities in the porosity can bias cell motion, thereby effectively inducing directed motion and giving rise to non-uniform cell density distributions. Due to its phenomenological similarity to other forms of directed cell motion [11], we frame this purely geometric effect as *porotaxis*. Our work establishes a direct link between the geometrical properties of tissue structures and cell migration modes, offering new insights into their interplay across diverse biological processes.

## II. MODEL

### A. Cell shape dynamics

To model cell motility within porous environments, we use the cellular Potts model (CPM) which accounts for cell shape dynamics. CPM is a widely used grid-based approach to describe various biological systems across multiple spatial and temporal scales. Within this framework, each lattice site is assigned a number (cell index). A single cell is then defined as the contiguous set of lattice sites with the same cell index. Cell properties and interactions between cells and their environment can be incorporated via the *Hamiltonian ℋ*, which is an energy-like quantity that governs the dynamics of the system. Since the framework is inherently discrete, additional layers of description, such as extracellular reaction-diffusion processes described by partial differential equations, can be readily integrated. For a comprehensive review on the CPM as a computational tool and its broad spectrum of applications, we refer to Refs. [42, 43]. We here choose a widely used form for the Hamiltonian defined by

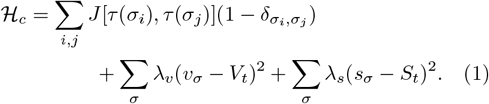

The first term on the right hand side in Eq. 1 accounts for cell adhesion between different cells *σ*_*i*_ of different cell types *τ* (*σ*_*i*_) with the elements of the interaction matrix **J** denoting the contact energies between cell type interfaces. The second term accounts for the fixed size of a cell and penalizes deviations of the actual size *v*_*σ*_ from the target cell volume *V*_*t*_. Analogously, the last term penalizes deviations of the cell surface *s*_*σ*_ from a target value *S*_*t*_. Taken together, the last two terms constitute the elastic energy of a cell, and the parameters *λ*_*v*_ and *λ*_*s*_, which define the strength at which deviations are penalized, may therefore be interpreted as stiffness parameters. The dynamics specified by Eq. (1) evolves in time via the *Metropolis-Hasting algorithm*, a Monte Carlo method that ensures that the state of the system converges to the Boltzmann distribution. Throughout this work, lengths are quantified in units of the CPM lattice size and time in units of Monte Carlo steps (MCS). Moreover, to mitigate bias from initial values, we incorporate a *burn-in phase* in the Monte Carlo algorithm, only using simulation data for downstream analysis after 1000 MCS. The values of model parameters are provided in Table I.

**TABLE I.** Model parameters. Lengths are given in units of CPM lattice size and energy is expressed in units of *k*_*B*_*T*.

| Parameter | Symbol | Value |
| --- | --- | --- |
| Cell-medium adhesion | $J_{cm}$ | 0 |
| Cell-cell adhesion | $J_{cc}$ | 10 |
| Cell-fiber adhesion | $J_{cf}$ | 0 |
| Temperature (Monte Carlo) | T | 2 |
| Volume constraint | $\lambda_v$ | 1 |
| Surface constraint | $\lambda_s$ | 1.5 |
| Motility bias | $\mu$ | 3 |
| Target cell volume | $V_t$ | 2500 |
| Target cell surface | $S_t$ | 980 |
| Rotational diffusion | $D_r$ | 1/600 |
| Total lattice sites | $L^3$ | 400<sup>3</sup> |
| Fiber length | $L_f$ | 100 |
| Fiber radius | $R_f$ | 3 |

### B. Cell motility

The Hamiltonian in Eq. (1) specifies cell shape and contact energies, but does not yet describe active cell motion. Although it is possible to integrate many biochemical and mechanical details of cell motility into a computational model [44–46], this usually involves high computational costs, which limits the broad applicability.

Here, we follow previous work [32, 33, 35] and model cell motility by active Brownian motion (ABM) [4, 5, 7]. To this end, each cell is assigned a polarity vector 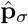 of unit length, which evolves over time according to the coupled system of stochastic differential equations (SDEs) defined by

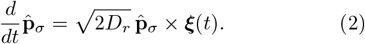

Equation (2) defines rotational diffusion of the 3D polarity vector 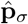 with a rate given by the rotational diffusion coefficient *D*_*r*_. The vector ***ξ***(*t*) describes uncorrelated Gaussian white noise with mean zero ⟨*ξ*_*i*_⟩ = 0 and variance ⟨*ξ*_*i*_(*t*) *ξ*_*j*_(*t*^*′*^)⟩ = *δ*_*ij*_*δ*(*t − t*^*′*^), and the cross product × denotes a Stratonovich product. The polarity vector thus mimics spontaneous cell polarization due to actin polymerization and accumulation, preferring cell protrusions and hence active motion in direction of 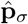 (Fig. 1(a)). Using spherical coordinates, the dynamics of 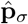 can be expressed by two SDEs for the azimuthal *θ* and polar angle *φ* (Fig. 1(b))

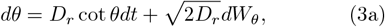

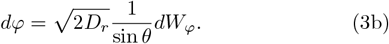

**FIG. 1.**
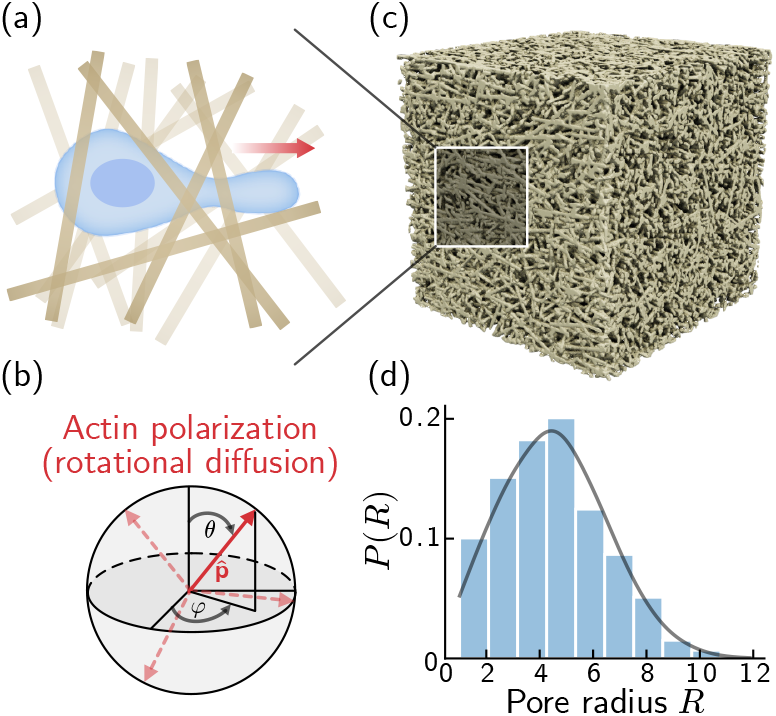
Model description. (a) Schematic illustration of cell motion through porous environments. A cell (depicted in blue) migrates through the void space (pores) of the medium. To move forward, a pseudopod forms in the direction dictated by the polarity vector (red arrow), and the cell’s body must deform to squeeze through small pores. (b) Illustration of the polarity vector 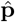 (red solid arrow) which mimics actin polarization and performs rotational diffusion on the unit sphere. The dashed red arrows visualize the polarity vector at different time points. (c) Exemplary disordered fibrous porous media as used in our simulations. The fiber radii and lengths are fixed to *R*_*f*_ = 3 and length *L*_*f*_ = 100, respectively (in units of CPM lattice size). The total length in all three spatial dimensions is set to *L* = 400. (d) Pore size distribution for the geometry shown in (c), showing that pore spaces are non-uniformly distributed. The characteristic pore size is given by the mean of the distribution 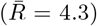 with the porosity here given by *ϕ* = 0.6.

Here, *dW* = *ξdt* denotes a Wiener process (Brownian motion), and the noise is interpreted in the Itô sense. Note that, in contrast to Eq. (2), *dW*_*θ*_ and *dW*_*φ*_ are independent Brownian motions (Appendix C). We utilize Eqs. (3) in our simulations because this form allows for more efficient numerical integration than Eq. (2).

To couple the dynamics of the polarity vector to the CPM framework, we add an additional term to the Hamiltonian in Eq. (1), i.e.,

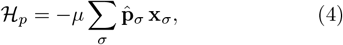

which introduces a bias for lattice updates in the direction of 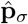. Here, the vector **x**_*σ*_ denotes the cell displacement induced by the update scheme in Eq. (1), with the parameter *µ* determining how strongly this bias is enforced. As a result, the combination of Eqs. (1),(3), and (4) leads to cell protrusion and subsequent movement in the direction 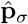.

As in active matter theory [28, 32, 33, 35, 47, 48], the cell motility model specified above is well-known to produce a persistent random walk (PRW) in free space (Movie 1), characterized by ballistic motion at short time scales and a transition to normal diffusion at larger time scales. The transition point between these two regimes is given by the persistence time *τ*_*p*_, which is related to the rotational diffusion coefficient *τ*_*p*_ = 1*/*2*D*_*r*_. Importantly and in contrast to (microscopic) active matter systems, for which agents are often assumed to be point-like, disk-shaped, or spherical [28, 40, 49, 50], our approach explicitly accounts for dynamic cell body deformations. This feature of our model is crucial because it captures effects of confinement, such as when cells must deform their bodies to squeeze through small pores or channels (Fig. 1(a)).

### C. Porous geometry

To investigate how structurally complex environments affect cell motility, we generate simulation geometries that resemble the disordered porous architecture of collagen-based ECM. Mimicking the disordered fibrous porous geometry, we generate a three-dimensional domain of total size *L* = 400 in all directions, consisting of randomly oriented and spatially distributed overlapping cylindrical fibers (Fig. 1(c)). Each fiber is assigned a fixed radius *R*_*f*_ = 3 and length *L*_*f*_ = 100 with all units given in CPM lattice sizes. Fibers are added iteratively until the desired porosity

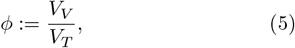

defined as the ratio of void volume *V*_*V*_ and total volume *V*_*T*_, is reached (Appendix D). The porosity can equivalently be expressed in terms of the reduced density 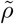

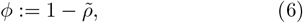

which is defined as the ratio between bulk density and collagen density. In experimental setups, 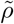 is usually the directly controllable parameter [14, 18, 51]. The obtained structure typically exhibits a non-uniform distribution of pore radii *R*, while remaining isotropic, i.e. the statistical properties of the fiber arrangement are similar in all spatial directions (Fig. 1(d)). The pore size distribution depends on the porosity *ϕ*: lower porosity results in smaller, more confined pores, while higher porosity leads to larger and more interconnected void spaces. Hence, the porosity *ϕ* influences the degree of confinement experienced by migrating cells within the medium.

These porous geometries are incorporated into the CPM simulation as binary voxel images, for which, for simplicity, the fibers are treated as rigid and non-deforming obstacles that constrain cell movement (Fig. 1(c)). Periodic boundary conditions are assumed in all spatial directions to mimic an unbounded environment.

## III. RESULTS

### A. Porous environments lead to transient diffusive regimes

To investigate how disordered porous environments affect cell migration, we simulate these dynamics within synthetic structures with varying porosities using the computational approach described in detail in Sec. II. To this end, *N* = 200 cells are randomly initialized within the synthetic porous environments and cell dynamics are simulated and tracked over a long time range of up to *T* = 10^7^ Monte Carlo time steps (MCS). For each cell, we then calculate the mean squared displacement (MSD) over time by

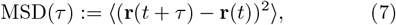

where **r**(*t*) denotes the position of the cell’s center of mass, *τ* the lag time, and ⟨·⟩ the ensemble and time average across all cells. Hereby, our target cell volume is set to *V*_*t*_ = 2500, which corresponds to a cell radius of *R*_cell_ = 8.4 (equivalent to the radius of a sphere with the same volume). The choice of the cell volume constrains the range of relevant porosities. If the porosity is too large, it would result in pore spaces larger than the cell radius, which is unphysical within our framework. Conversely, if the porosity is too small, it would completely block any cell movement. Therefore, based on our parameter choice, we perform simulations across eight relevant geometries with varying porosity within the interval *ϕ* ∈ [0.52, 0.70]. With this choice, the cell radius is approximately twice as large as the mean pore radius of the least dense medium *ϕ* = 0.7, i.e., 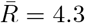 (Fig. 1(d)), requiring cells to deform their bodies when moving through the pore space.

Our results show that cell motility within porous environments deviates from persistent motion and exhibits transient motility regimes (Fig. 2(a)). At short times, the MSD displays superdiffusion with scaling exponent *β >* 1 and MSD(*τ*) ∼*τ* ^1.7^. At intermediate times, spanning up to three orders of magnitude, cells exhibit subdiffusive motion, with the scaling exponent *β <* 1 of the MSD decreasing as confinement increases (Fig. 2(b)). Interestingly, the superdiffusion exponent is independent of porosity, which can be seen by rescaling the lag time and MSD using the cross-over time between superdiffusive and subdiffusive motion (Figs. 2(a) and 2(b)). At large time scales, the MSD transitions again from subdiffusion to normal (Brownian) diffusion, except for porosities below a critical point around *ϕ*_*c*_ = 0.6, where subdiffusive motion persists.

**FIG. 2.**
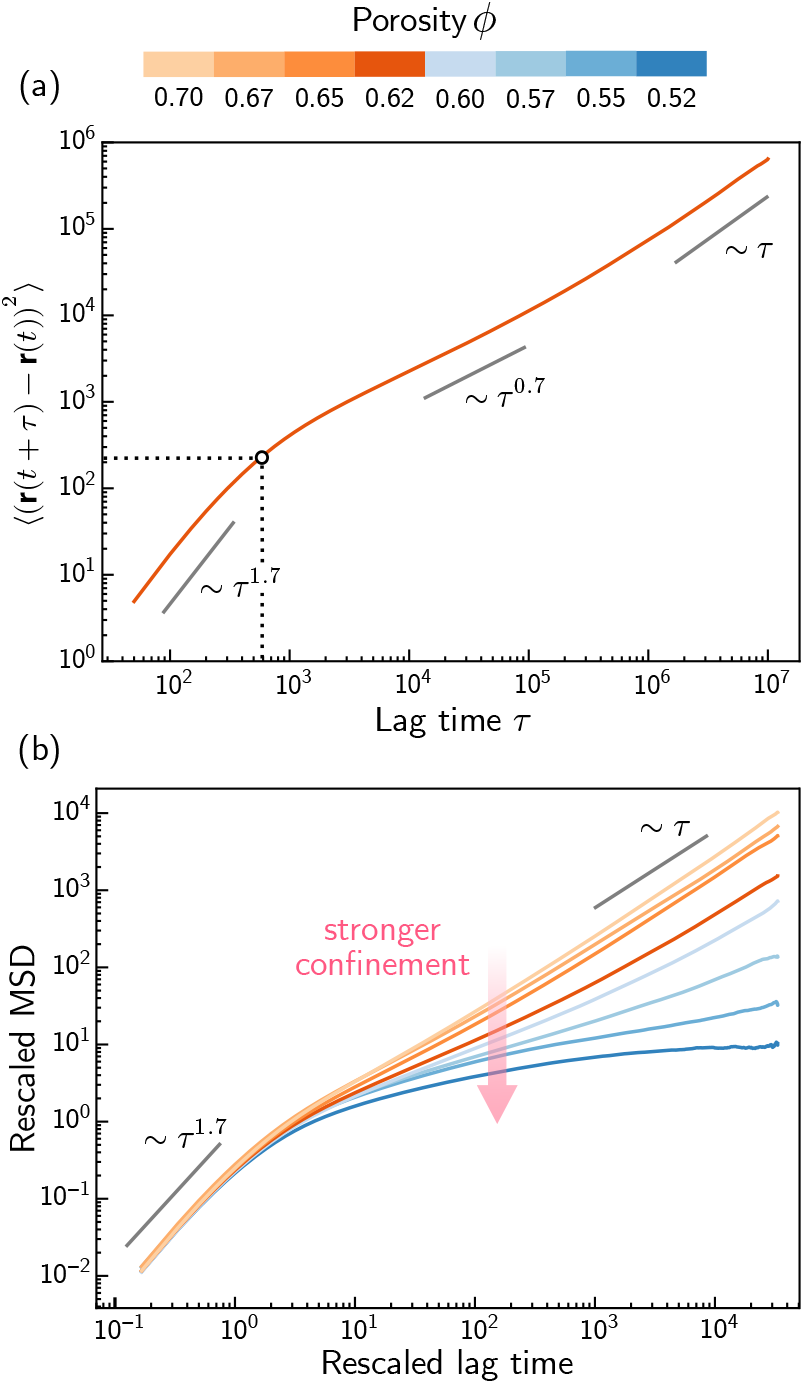
Mean-squared displacement across various porosities. (a) MSD of a single cell in a porous environment with porosity *ϕ* = 0.62, showing that cell motility exhibits transient regimes of diffusion. (b) Ensemble-averaged MSD for geometries with different porosities. For each curve, the MSD and lag time have been rescaled by the cross-over time from superdiffusion to subdiffusion (dashed lines in (a)). This shows that the superdiffusion exponent is independent of the porosity. The long-time dynamics, however, is sensitive to the porosity and transitions from normal diffusion to subdiffusion around the critical porosity *ϕ*_*c*_ = 0.6.

Taken together, our findings suggest that cell motility in random porous media resembles ‘hopping between traps’ rather than persistent motion: Cells trapped in small pores must explore the pore space through protrusions until they discover a suitable path to adjacent pores (Movie 2) leading to heterogeneous movement patterns characterized by fast ‘hops’ and slow exploration phases when ‘trapped’. Depending on pore size and its connectivity to neighboring pores, the trapping time can be significantly prolonged, which aligns with the observed dependency of the subdiffusion time range on the porosity of the environment observed in the MSD (Fig. 2(b)). In the following, we will show that the different diffusion regimes can be fully characterized by statistical properties of the hopping and trapping dynamics.

### B. Pore space disorder shapes the statistical properties of cell motility

Random walks and transport processes in disordered media can be well described by continuous-time random walk models (CTRWs) [26, 27, 52]. Here, the random walk is governed by a master equation for the probability density *P* (**r**, *t*) to find a walker at position **r** and time *t*. The form of the master equation and the resulting dynamics generally depend on two key factors: the run distance of a walker, characterized by a probability density function (PDF) Λ(**r**) defining the ‘jump length distribution’, and the time a walker waits before it makes the next ‘jump’, drawn from another PDF *ψ*(*t*), characterizing the waiting time distribution. Based on assumptions about the properties of Λ(**r**) and *ψ*(*t*), CTRWs predict different regimes of diffusion, including anomalous diffusion and Lévy walks [26, 27]. For an excellent and comprehensive review on CTRW models, we refer the reader to the article by Zaburdaev et al. [27].

The points above suggest that the statistics of hopping length *ℓ*_*h*_ - the distance a cell travels before getting trapped - and trapping duration *τ*_*t*_ - the time a cell remains in a pore before hopping again - may explain the different diffusion regimes observed in the MSD. The distributions of *ℓ*_*h*_ and *τ*_*t*_ can be determined from cell tra-jectories, as explained below. We first calculate the cell displacement

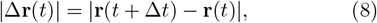

for fixed time increments of Δ*t* = 50. Multiplied by 1*/*Δ*t*, Eq. (8) would correspond to the instantaneous cell velocity. The displacement of a single cell typically shows intermittent phases of hopping and trapping (Appendix B). Hops are characterized by sudden changes in |Δ**r**(*t*)| over time, while traps are indicated by little to no change in |Δ**r**(*t*)|. By this, we are able to segregate trajectories into time intervals during which cells are either trapped or actively hopping. In order to account for stochastic effects as cell displacement is calculated based on variations in the center of mass of a cell, we classify trapping as time intervals where the cell displacement stays below a threshold value of |Δ**r**(*t*)| *<* 4, and hops as time intervals above this threshold (Appendix B). The threshold is chosen to be in the order of the typical pore size, such that the motion of the cell’s center of mass within a pore is classified as trapping. However, our conclusions do not depend on the specific value of the threshold. To proceed, we first determine the hop length and trapping times for each cell separately, and then combine these to obtain the ensemble distributions. For numerical convenience, we calculate survival probabilities (i.e., complementary cumulative distribution functions (CCDF)) instead of probability densities.

#### 1. Hopping length distribution

As shown in Fig. 3(a), we find that the distribution of hop lengths follows an exponential distribution. The characteristic decay length *λ* depends on the porosity of the environment, and decreases with smaller porosities, as one would intuitively expect. One question that arises here is whether the hop length distribution is guided by the microstructure of the porous environment. In general, the pore sizes and conduits that connect different pores determine the number and shape of percolating paths, thereby defining the transport properties in porous media. An important quantity in this context is the *chord-length distribution*, which describes the distribution of straight paths of length *ℓ*_*c*_ that fit into the pore space [53]. Using binarized voxel images of the simulated geometries, we determine the chord-length distribution and compare it to the hop length distribution (Fig. 3(a)). We find that the chord-length distribution aligns with the hop length distribution, strongly suggesting that cell movement is guided by percolating paths within the porous medium [52, 54].

**FIG. 3.**
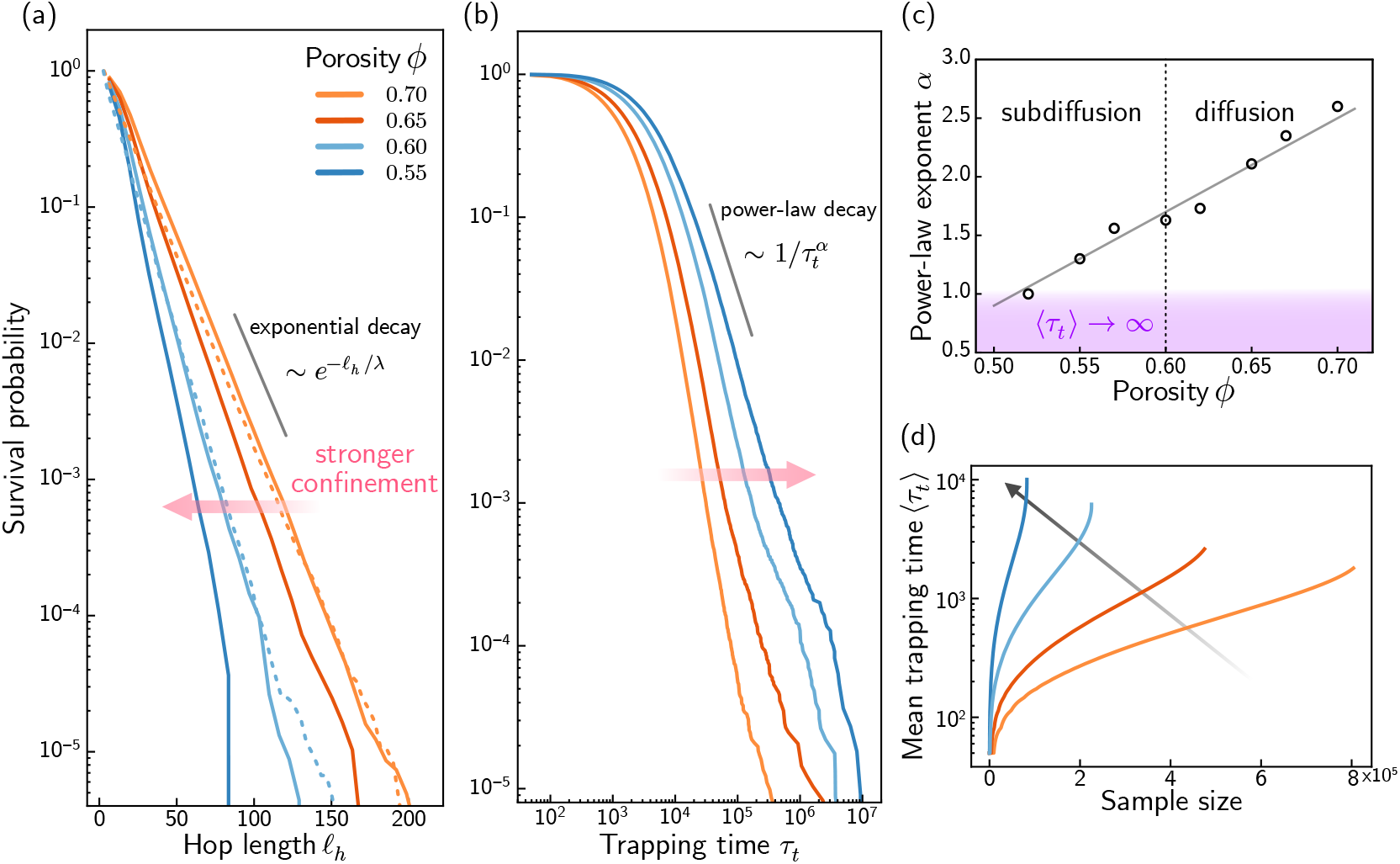
Statistical properties of cell motility in disordered porous environments. (a) The hop length distribution for different porosities (solid lines) follows an exponential distribution with the characteristic hop length *λ* decreasing with lower porosities. The hop length distribution corresponds to the chord length distribution, i.e., the characteristic distances between different pores (dashed lines, for clarity only shown for two porosities). (b) The distribution of trapping times exhibits a power-law tail that spans approximately three orders of magnitude. Lower porosities shift the tail towards longer trapping times, eventually causing the mean trapping time to diverge. (c) The value of the power-law exponent *α* for the trapping time depends weakly on the porosity. For a perfect power-law, the purple shaded region indicates the point for *α* below which the mean value of the distribution diverges. However, we observe a transition from diffusion to subdiffusion around *ϕ*_*c*_ = 0.6, corresponding to *α ≈* 1.5. (d) Mean trapping time for different sample sizes derived from the order statistics of the distributions shown in (b). The sample average (mean trapping time, ⟨*τ*_*t*_⟩) increases as the sample size grows. This behaviour is one hallmark of distributions with heavy tails, as it shows how outliers significantly impact the mean. Moreover, small changes in the power-law exponent *α* (cf. (c)) can lead to significant variations in the mean trapping time, eventually leading to divergence.

#### 2. Trapping time distribution

The trapping times are broadly distributed, covering up to seven orders of magnitude indicating that cells could be trapped within pores throughout the whole time period of the simulation, i.e., *T* = 10^7^. (Fig. 3(b)). Strikingly, the tail of the distribution shows a power-law decay 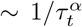 over approximately three orders of magnitude. The heavy-tail of the distribution has important implications for the dynamics: in particular, the first moment of power-law distributions only exists if *α >* 1, while it diverges for *α* ≤ 1 (note, however, that the distributions here are not perfect power-laws). Upon inspecting the trapping time distributions, we observe that the exponent *α* decreases weakly with smaller porosities (Fig. 3(c)), which corresponds to a substantial increase in the mean trapping time ⟨*τ*_*t*_⟩ (Fig. 3(d)).

#### 3. Implications from CTRW models within porous environments

The distributions of hopping lengths and trapping times, as characterized above, emphasize that cell motil-ity dynamics within porous environments can be interpreted within the framework of CTRW models. The exponential form of the hopping length distribution implies that spatial displacements are well-behaved and with a characteristic length scale set by the pore geometry. In particular, both, the first and second moment of the hop length distributions are finite. In contrast, the trapping time distributions are heavy-tailed, and the first moment, i.e. the mean trapping time ⟨*τ*_*t*_⟩, diverges for small porosities. In the CTRW framework, such heavy-tailed waiting times with diverging first moment are known to give rise to subdiffusion [26, 27]. This is consistent with our simulations, which show a transition from normal diffusion at high porosities to subdiffusive behaviour at low porosities (Figs. 2 and 3(c)). Importantly, if the mean trapping time is finite, the dynamics recovers normal diffusion at long time scales, regardless of the specific details of the hopping length distribution (as long as the moments exist). This is due to the central limit theo-rem, which ensures that the sum of independent random variables, drawn from any distribution with finite mean and variance, converges to a normal distribution [26, 27]. However, since the waiting times are also broadly distributed in this case, the dynamics at intermediate time scales can be subdiffusive due to trapping [26]. This is, again, consistent with our simulation results which show transient subdiffusive regimes for *ϕ > ϕ*_*c*_ (Figs. 2 and 3(c)).

Overall, the combination of exponentially distributed hopping lengths and heavy-tailed trapping times within porous environments aligns well with CTRW models and provides a mechanistic explanation for the different diffusion regimes observed in our simulations. Notably, the porosity is the relevant control parameter that determines the transition from normal to anomalous diffusion.

### C. The microstructure dictates the effective diffusion coefficient within porous environments

#### 1. Large-scale diffusion coefficient

For porosities *ϕ > ϕ*_*c*_, cell motility is well described by normal diffusion. This is particularly valid for coarse-grained dynamics at length scales comparable to ⟨*ℓ*_*h*_⟩ and time scales larger than ⟨*τ*_*t*_⟩. At these scales, cell motility in porous environments may be envisioned as a random walk on a disordered lattice, with mean squared length of a jump set by 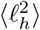 and mean hopping rate to neighboring sites given by ⟨*τ*_*t*_⟩^*−*1^. Assuming that the dynamics is isotropic in all spatial directions, it is straightforward to show that the effective diffusion coefficient 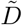 at large times is given by [26]

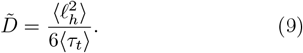

As shown in Fig. 4(b), the value determined from Eq. 9 is a reliable estimator for the effective diffusion coefficient, which we confirm through comparison with the value 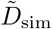 extracted from the MSD in our simulations, i.e.,

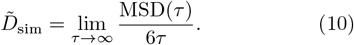

**FIG. 4.**
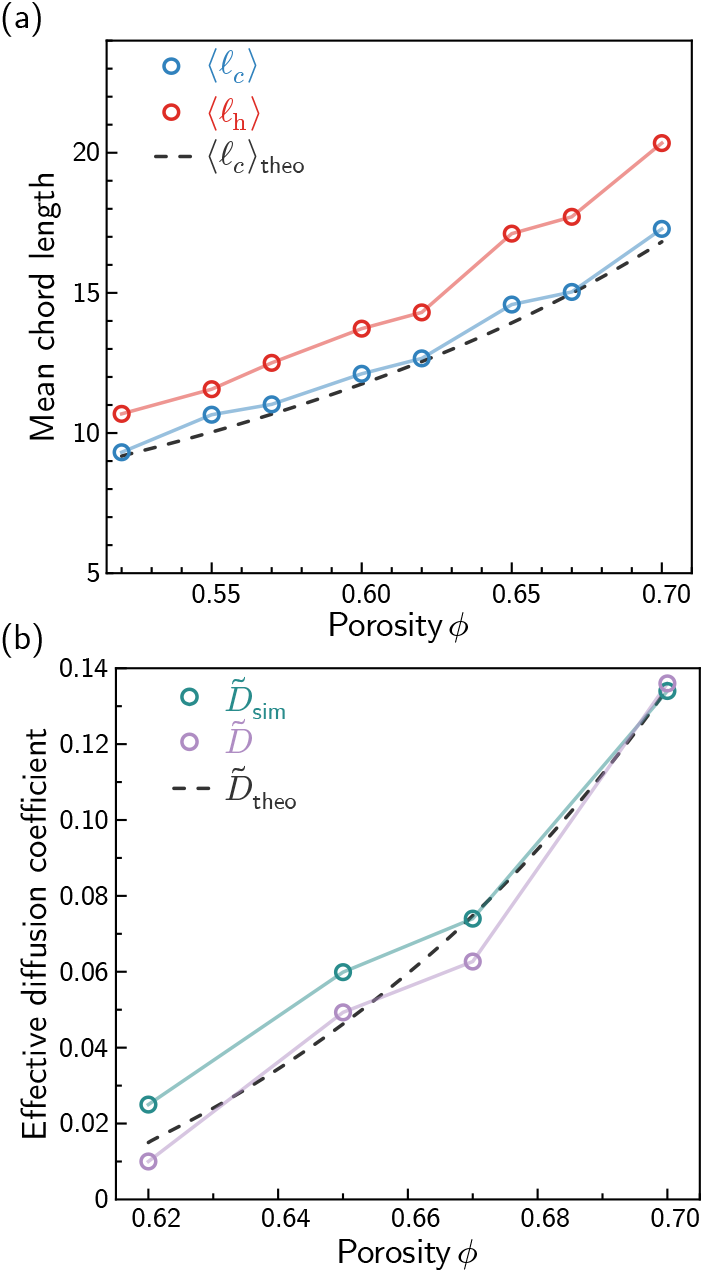
The microstructure of the porous medium governs geometric characteristics and large scale transport properties. The mean hop length (⟨*ℓ*_*h*_⟩, solid red line), mean chord length (⟨*ℓ*_*h*_⟩, solid blue line), and the mean chord length as theoretically predicted by Eq. 15 (⟨*ℓ*_*h*_⟩ _theo_, dashed black line) for various porosities. (b) The effective diffusion coefficient obtained from the MSD (solid teal line) agrees fairly well with the values calculated from the simple expression in Eq. 9 (solid purple line). The theoretical prediction dependent on characteristic values of the microstructure from Eq. 21 (dashed black line) aligns well with these values.

#### 2. Mean chord length solely depends on fiber density and radius

In order to relate 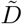 as defined by Eq. 9 to geometric properties of the porous environment, we leverage on the observation that hopping lengths are guided by the chords of the porous medium (Figs. 3(a) and 4(a)). This allows us to approximate the mean hop length by the mean chord length ⟨*ℓ*_*h*_⟩ *≈* ⟨*ℓ*_*c*_⟩ . To determine how ⟨*ℓ*_*c*_⟩ further relates to the porosity of the environment, we borrow ideas from geometric probability and kinetic theory [55–57]. First, the porosity *ϕ* can be interpreted as the probability that a random point lies within the void space, i.e., the point is not covered by any cylindrical fibers. For non-overlapping cylindrical fibers, this probability follows directly from the definition in Eq. 5 with

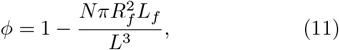

where *N* denotes the total number of fibers, *L* the maximal dimension of the simulated cube, and *L*_*f*_ and *R*_*f*_ the length and radius of a fiber, respectively (cf. Sec II C). However, for overlapping fibers, as in the case here, Eq. 11 underestimates the probability, since overlaps reduce the total fraction of space covered by fibers. To generalize Eq. 11, consider *N* randomly distributed points within the total volume *V*, where each point corresponds to the midpoint of a randomly oriented cylindrical fiber. We then ask for the probability *P* (*k*) of finding *k* midpoints within a test volume of size 𝒱. For sufficiently small test volumes 𝒱, *P* (*k*) follows a Poisson distribution with

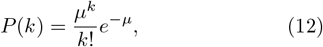

where the parameter *µ* is given by *µ* = *n𝒱*, and *n* denotes the number density of midpoints, i.e., *n* = *N/L*^3^. The porosity then follows from Eq. 12 by determing the probability of finding no point, *k* = 0, within the excluded volume 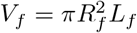 (i.e., the volume covered by fibers). Hence, we obtain

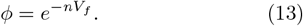

Note that in the dilute limit *nV*_*f*_ ≪ 1, where fiber overlaps are rare, Eq. 13 reduces to Eq. 11 for non-overlapping fibers. Next, we interpret the mean chord length as the *mean free path* of a particle beam traveling through the porous medium. In this picture, the average distance a particle travels before being stopped or scattered by obstacles, ⟨*ℓ*_*c*_⟩, is theoretically related to the number density of obstacles *n* and the effective cross-sectional collision area *σ*,

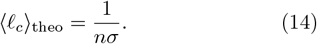

For randomly oriented cylindrical fibers, and assuming that *L*_*f*_ ≫ *R*_*f*_, the effective cross-section area is given by *σ* = *πR*_*f*_ *L*_*f*_ */*2. Heuristically, the factor of two in *σ* reflects that a particle can only ‘observe’ one half of the cylindrical obstacle. Finally, combining Eqs. 13 and 14, we conclude that

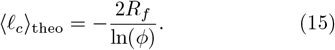

Interestingly, the mean chord length does not explicitly depend on fiber length *L*_*f*_ . This is because the fiber length contributes an additive factor of order ∼1*/L*_*f*_ to the mean chord length, which is subleading compared to the term in Eq. 15. Comparison of the predicted value from Eq. 15 and the numerically determined mean chord length shows excellent agreement (Fig. 4(a)).

Exploiting the fact that chord lengths are exponentially distributed, i.e. using that 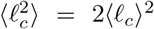 and ⟨*ℓ*_*h*_⟩ *≈* ⟨*ℓ*_*c*_⟩, we can combine Eqs. 9 and 15 to arrive at the following expression for the effective diffusion coefficient

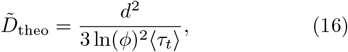

with *d* ≡ 2*R*_*f*_ denoting the fiber diameter.

#### 3. Mean trapping time is governed by entropic traps

Analogously, it remains to be determined how the mean trapping time ⟨*τ*_*t*_⟩ in Eq. (16) is linked to the microstructure of the porous environment. To this end, we utilize the concept of *entropic traps*, a model that has been used to explain power-law trapping time distributions across diverse physical systems, including confined DNA molecules [58], motile bacteria [54], and active polymers in porous media [52]. The foundations of this concept were laid by Robert Zwanzig in his work on ‘diffusion past an entropy barrier’ [59].The basic idea is to determine the free energy difference Δ*F* required by a cell to escape a pore (‘trap’). This energy barrier is associated to all possible conformational states of a cell that result in trapping, Ω_*t*_, and all states that enable the cell to escape the trap, Ω_*e*_, i.e.,

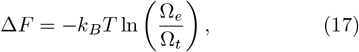

with *k*_*B*_ denoting the Boltzmann constant and *T* the temperature of the system. In general, the conformational states and, consequently, Δ*F* are governed by intricate microscopic details, such as pore sizes and pore connectivity, pore throat dimensions, cell size and shape, cell mechanics, and adhesion-mediated interactions with pore surfaces. As it is challenging to rigorously incoporate all these microscopic details to Eq. (17), we instead propose a phenomenological approach based on scaling arguments. In short, we assume for *ϕ > ϕ*_*c*_ that the average number of states obeys a scaling law of the form

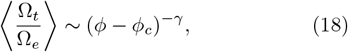

where, in analogy to percolation theory, the value of the positive scaling exponent *γ* generally depends on model assumptions and space dimension [60], and we here set *γ* = 1 for simplicity. Next, we assume that the mean trapping time follows an Arrhenius law,

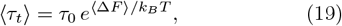

such that large energy barriers (i.e., small pores) lead to exponentially growing mean trapping times. The parameter *τ*_0_ in Eq. (19) denotes the characteristic *attempt time* to escape a trap, and generally depends on microscopic parameters of the system [61]. Combining Eqs. (17)–(19)^1^, we find that

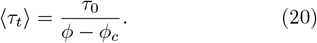

Finally, merging Eqs. (16) and (20) we arrive at an expression for the effective diffusion coefficient with regard to the microstructure of the environment

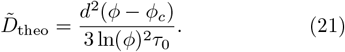

Equation (21) coarse-grains model details at small scales into two free parameters, *ϕ*_*c*_ and *τ*_0_. Using the fact that here *ϕ*_*c*_ = 0.6 (cf. Fig 3(c)), we fit Eq. (20) to the mean values of the trapping time distribution obtained from our simulations, resulting in *τ*_0_ ≈ 70. As shown in Fig. 4(b), the theoretically derived effective diffusion coefficient dependent on characteristics of the porous geometry (Eq. 21) agrees fairly well with the values obtained from the simulation and the simple expression in Eq. 9.

### D. Porotaxis in heterogeneous porous media

#### 1. Porosity gradients effectively induce directed cell motility

The ECM is rarely uniform in its properties and spatial distribution. Multiple factors may cause spatial inhomogeneities, including interactions with the cells themselves as well as pathological conditions, such as viral infections and cancer [7, 62, 63]. This led us to the question of how spatial heterogeneities in porosity affect the spatial distribution of cells in porous environments. One hallmark of confinement is the slowing down of cell motility, arising here from trapping events in the porous environment. Indeed, slowing down depends strictly on the microstructure and is reflected by a decrease of the large scale effective diffusion coefficient with decreasing porosity (cf. Fig. 4(b)). As a consequence, if porosity varies in space, one may expect that cells spend more time in regions of higher confinement due to trapping and the overall reduced mobility of cells, suggesting non-uniform spatial distribution of cells.

To test this, we perform simulations in heterogeneous geometries with various spatial porosity gradients. We construct 3D simulation geometries in which the porosity changes along the z-axis. Simulating the dynamics of *N* = 200 cells followed over *T* = 10^6^ time steps we then examine the stationary density distribution of cell positions along the z-axis for all cells at times *t > T*_*s*_ = 10^5^.

Our simulations show that spatial heterogeneities induce non-uniform cell distributions, with cells clearly accumulating in regions of lower porosity where confinement is more pronounced (Fig. 5 and Movie 3). This effect persists irrespective of the specific form of the porosity gradient, i.e., considering either abrupt (Fig. 5(a)) or gradual changes in porosity (Fig. 5(b)). Thus, the stationary density increases toward regions of lower porosities. In other words, our findings suggest an *effective di-rected cell motion* downstream of porosity gradients, rem-iniscent of directed cell migration observed under chemo-tactic stimulation (chemotaxis). Since the effective directed motion here is attributed to spatial gradients in pore confinement, we refer to this mechanism as *poro-taxis*.

**FIG. 5.**
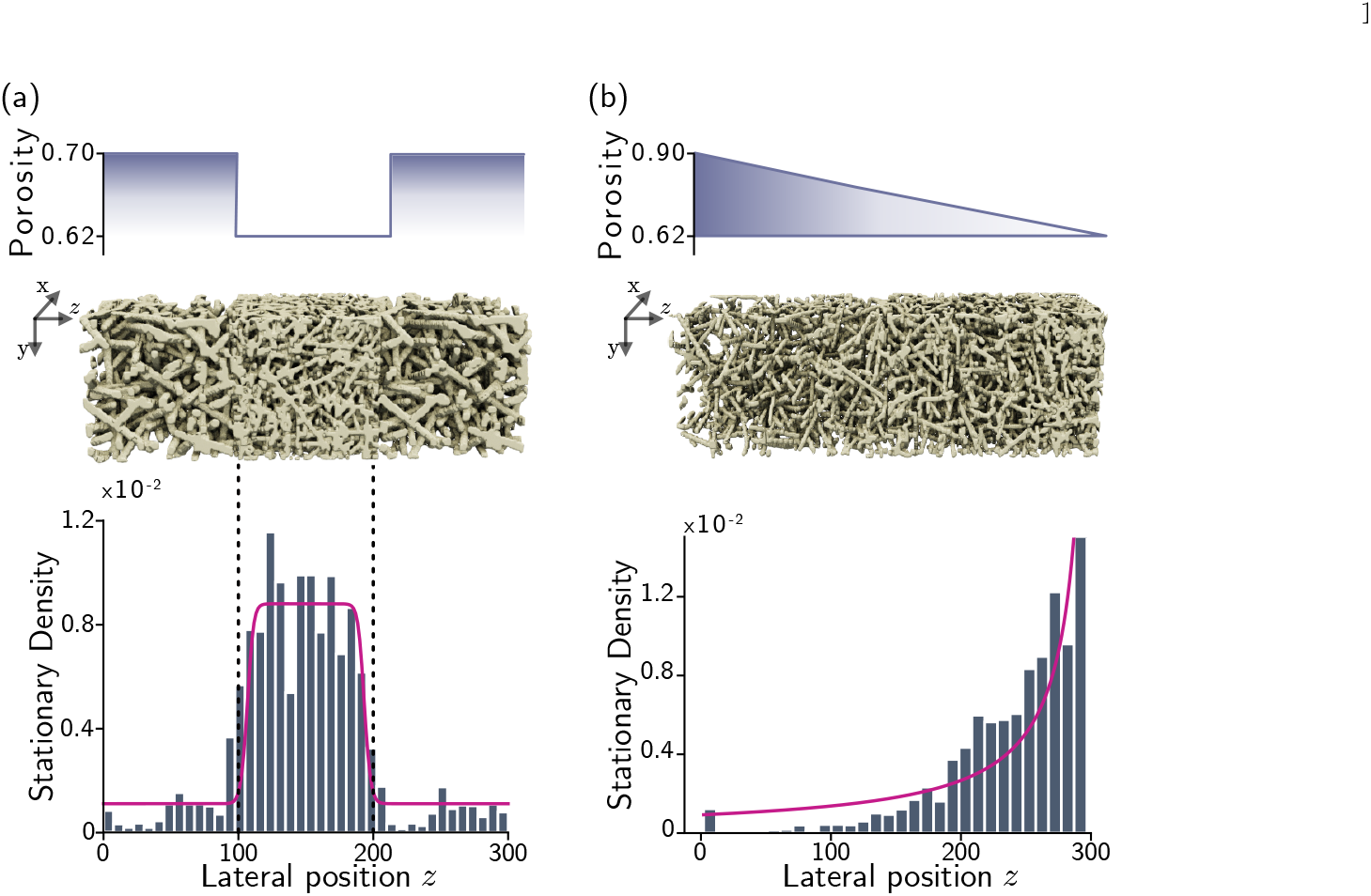
Porotaxis in heterogeneous porous media. Stationary cell density distribution along the lateral position (z-axis) in heterogeneous 3D environments with (*x, y, z*)-dimensions 100 *×* 100 *×* 300, assuming (a) an abrupt change in porosity from *ϕ*_+_ = 0.7 to *ϕ*_*−*_ = 0.62, and (b) a linear decrease in porosity along the lateral position with *ϕ*(*z*) = *ϕ*_+_ *− z* (*ϕ*_+_ *− ϕ*_*−*_) */L*_*z*_ and *ϕ*_+_ = 0.9 and *ϕ*_*−*_ = 0.62. Periodic boundary conditions are assumed along the x- and y-axis, while no-flux boundary conditions are assumed along the z-axis to allow the system to reach a steady state. Initially, *N* = 200 cells are randomly distributed within the domain, corresponding to homogeneous cell density distributions. Cells are followed for *T* = 10^6^ simulated time steps. Final stationary density distributions along the z-axis are determined by the histograms of the z-component *r*_*z*_ of the position vector **r**(*t*) for all cells at times *t > T*_*s*_ = 10^5^. In both geometries, the homogeneous initial condition is unstable and evolves into a non-uniform stationary density profile that reaches its peak in regions of low porosity (histogram in (a) and (b)). The stationary solution Eq. (24) (purple solid lines in (a) and (b)) aligns well with the simulation results.

#### 2. Inhomogeneous diffusion underlies porotaxis

Exploiting the fact that, at sufficiently large scales, the dynamics is governed by normal diffusion for *ϕ > ϕ*_*c*_, cell motility in heterogeneous porous environments may be viewed as a diffusion process with spatially varying diffusivity. Specifically, we can model the *z*-coordinate of cells using a Langevin equation with position-dependent diffusivity 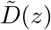,

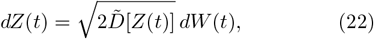

with the multiplicative noise interpreted in the Itô sense. Equation (22) is equivalent to a Fokker-Planck equation describing the time evolution of the probability density *p*(*z, t*) to find a cell at position *z*,

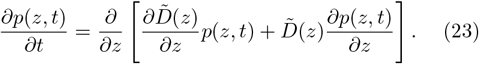

For closed systems with reflective boundary conditions *∂*_*z*_*p*(*z, t*) |_*z*=0_ = *∂*_*z*_*p*(*z, t*) |_*z*=*L*_ = 0, Eq. (23) admits the stationary solution

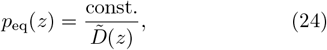

with the constant factor on the right-hand side determined by normalization of *p*_eq_(*z*). Equation (24) indicates that cells accumulate in regions of low porosity, for which 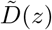 is reduced. From the equilibrium solution Eq. (24), we furthermore obtain an *effective potential U* (*z*) [64],

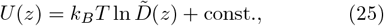

which emphasizes inhomogeneities in 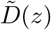 as the driving force underlying porotaxis. Extracting the values of 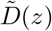 from the microstructure (cf. Sec. III C), we find that the stationary solution Eq. (24) matches our simulations fairly well (Fig. 5(a),(b)).

## IV. DISCUSSION

By combining computational modeling and theory, we showed how disordered porous environments control the mode of cell motility. While, heuristically, one would expect that confinement solely reduces the persistence time, as the characteristic time for unobstructed movement relates to the porosity, our results underscore that cell migration in porous media deviates from persistent motion and is best understood as a random walk in a disordered landscape of traps. Cell movement is governed by hops between pores, with the hopping rate set by the inverse of the average time a cell spends within a pore (i.e., being trapped).

Our results are consistent with, and complement, previous experimental studies on bacterial motility in porous hydrogels, where hopping and trapping have also been observed and characterized [54, 65, 66]. Strikingly, other biological systems also exhibit hopping and trapping dynamics in disordered environments, such as active polymers in granular porous media [52], as well as molecules within disordered nanoporous media [67]. Although microscopic details and motility differ between these systems and ours, the resulting dynamics share common features. Bacteria, for instance, typically exhibit a stiff spherocylindrical shape and migrate via run-and-tumble motion[47], whereas active polymers are characterized by elongated, fluctuating conformations and move by randomly switching between runs and reversals [52]. This is in contrast to our system, in which the cell’s direction of movement evolves continously over time due to rotational diffusion in ABM. Crucially, since we model the cell body as a deformable elastic object, cells are able to squeeze through pores smaller than their size – a process not possible for bacteria or active polymers. Despite these differences, the large scale dynamics is identical (above *ϕ*_*c*_) and governed by normal diffusion, where the effective diffusion coefficient comprises geometric and (coarse-grained) model specific details. This suggests that, at a coarse-grained level, microscopic details of cell migration may not be relevant. However, the onset to subdiffusion motion at large time scales for porosities *ϕ < ϕ*_*c*_ has not been observed for bacteria and active polymers in porous media. We attribute this difference to the elasticity of the cell body – a feature not present in bacterial cells and polymers.

By linking chord lengths and mean trapping times to the porosity, we have shown that the characteristic length and time scales of these ‘hopping and trapping’ dynamics are fully characterized by geometric properties of the microenvironment. This, in turn, enabled us to connect the large scale effective diffusion coefficient to these geometric parameters. In line with these findings, we have shown that spatially heterogeneous environments effectively induce directed motion of cells towards regions of lower porosity (i.e., porotaxis). On a mechanistic level, porotaxis results from inhomogeneous diffusion, characterized by an effective potential field, thereby leading to stationary non-uniform cell density distributions.

It is well known that several external factors, such as biochemical and mechanical cues, can control and guide cell motion[11]. Representative examples are diverse and include motion along gradients of chemokines (chemotaxis) [15, 19, 20, 68], ECM stiffness (durotaxis) [69], electrical fields (galvanotaxis) [2], ligand concentration (haptotaxis) [7], as well as topography (topotaxis) [40, 70] and curvature (curvotaxis) [71]. The effective directed motion, porotaxis, described in this work, shares the same phenomenology as the examples highlighted above, namely biased movement in response to spatial cues. However, the mechanism underlying porotaxis is substantially different from directed motion due to biochemical and mechanical signalling. For instance, chemotaxis is driven by a cascade of intricate biomolecular processes, involving sensing of ligands and ensuing intracellular information processing that feed back to cell motion [72, 73]. As such, chemotaxis is an active and versatile process that can be regulated in a multitude of ways, including by cells themselves (autochemotaxis) and environmental factors [7, 11]. In contrast, porotaxis is a purely geometric effect and emerges as a result of increased trapping events in regions of lower porosity. Since this effect is based on spatial variations in confinement, it may be controlled through ECM remodeling [7, 14] or by cells switching to a mesenchymal migration program (e.g., via mechanosensing) [74, 75]. However, the time scale of ECM remodeling is generally longer as compared to, for instance, chemokine signaling [11, 14], illustrating that porotaxis operates as a rather slower guidance mechanisms.

Notably, cancer cells and viral infections are known to remodel the ECM into denser, fibrotic regions [62, 76, 77], suggesting that they may exploit porotaxis to influence immune cell infiltration and invasion dynamics, and, thus, tissue organization and function. More broadly, one might suggest that porotaxis could interact with other guidance mechanisms. For example, during a viral infection, chemotactic gradients could direct immune cells toward sites of inflammation, while structural changes in the ECM may additionally support this process through porotaxis. This also applies to observed motility modes of T cells that are associated with improved antigen recognition capabilities and could be linked to structural elements based on our findings [17, 78]. Such synergy could provide an additional means of fine-tuning immune cell recruitment and the spatial organization of cells within complex tissue environments.

In conclusion, cell migration is shaped by a complex interplay between mechanical, biochemical, and environmental cues [3, 75, 79, 80]. Understanding how the structural properties of the ECM contribute to this interplay is therefore essential for disentangling the respective roles of molecular signaling and physical constraints during many biological processes. By combining computational modeling with theoretical analysis, we established a direct link between the geometric features of disordered environments and emergent motility dynamics. Our approach not only provides a conceptual framework for interpreting experimental observations, but also reveals how structural heterogeneity can control cell motion, opening new avenues to better understand diverse complex processes, including chronic inflammation, immune surveillance, and cancer invasion.

## ACKNOWLEDGMENTS

The authors would like to thank Jörn Starruß and Lutz Brusch for discussions and their helpful assistance with the implementation of the CPM model, and Isabella R. Graf for valuable feedback on the manuscript. This work was supported by the German Federal Ministry of Research, Technology and Space (BMFTR) within the Computational Life Science Initiative (EMUNE/031L0293E to F.G.). F.G. was additionally supported by the Hightech Agenda (HTA) Bavaria. L.W. and F.G. conceived the study and designed the computational experiments. L.W. developed the model, performed the simulations, and analysed the data. L.W. and F.G. interpreted the results and wrote the manuscript.

## Appendix A: CPM implementation and parameters

The CPM model has been implemented in *Morpheus* version 2.3.8, a freely available open-source software package [81]. The simulation used to generate all results in this work is available as an *XML* file on GitHub. This file can be opened directly from the graphical user interface of *Morpheus*. The model parameters used in this study are summarized in Table I.

## Appendix B: Hopping and trapping statistics from cell displacement

We use the instantaneous cell displacement Eq. (8) to determine hopping and trapping phases for each cell, as illustrated in Fig. 6. Before classification, the cell displacement is smoothed by using a moving average over the last four values for each time point. To determine hop lengths, we calculate the average cell velocity over all hopping phases and multiply this value by the duration of the hops (Fig. 6). Hop lengths and trapping times from all cells are pooled to obtain the distributions shown in Figs. 3(a) and 3(b).

**FIG. 6.**
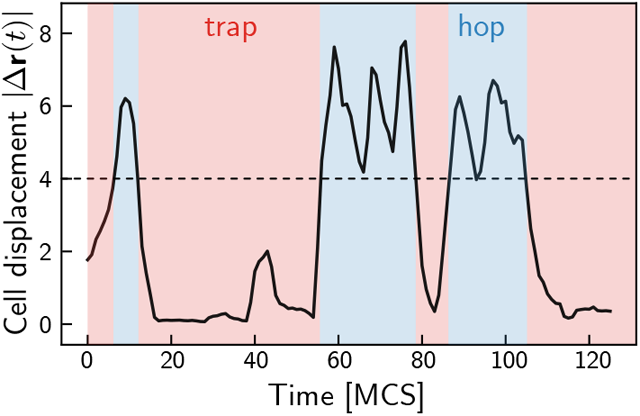
Representative cell displacement of a single cell (black solid line). Time intervals where the cell displacement stays below the threshold value |Δ**r**(*t*)| *<* 4 (dashed black line) are considered as traps (red shaded area), and time intervals where the displacement exceeds this threshold represent hops (blue shaded area).

## Appendix C: Brownian motion on the unit sphere

In this section, we provide details for the derivation of the SDEs describing the time evolution of the azimuthal and polar angle (see Eqs. (3)). Using spherical coordinates and dropping the cell index *σ* for clarity, we first rewrite the time evolution of the polarity vector as

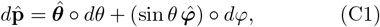

where we explicitly made use of the fact that the SDE is defined in the Stratonovich sense (as indicated by the Stratonovich product ∘). Here, 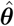 and 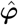 denote the orthogonal unit basis vectors in the directions *θ* and *φ*.

Expanding Eq. (2) in spherical coordinates

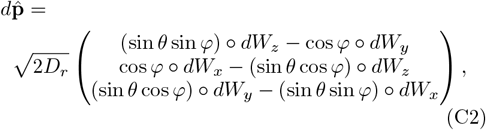

where *dW*_*i*_ = *ξ*_*i*_*dt* denotes the Wiener process along the spatial direction *i* = {*x, y, z*}, and recalling that

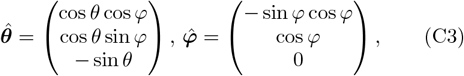

one can determine expressions for *dθ* and *dφ* by comparing Eqs. (C1) and (C2)

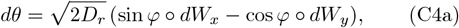

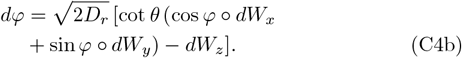

Defining *d***X** := [*dθ, dφ*]^*T*^, the above equations can be alternatively written in matrix form

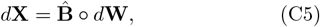

with the elements of the matrix *B*_*ij*_ given by the respective terms in Eqs. (C4a) and (C4b), and here *i* ∈ {*θ, φ*} whereas *j* ∈ {*x, y, z*}. Introducing the drift term *A*_*i*_ = *B*_*kj*_*∂*_*k*_*B*_*ij*_, where *∂*_*k*_ denotes the partial derivative with respect to *θ* and *φ*, respectively, and using Einstein summation, we convert Eqs. (C4a) and (C4b) to the equivalent Itô form [82]

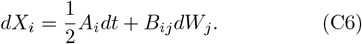

Expanding each term explicitly in Eq. (C6), we find

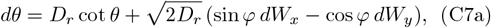

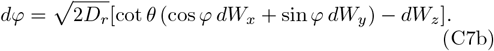

Noting that

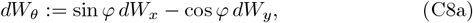

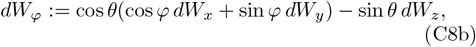

are two independent Wiener processes, one can rewrite Eqs. (C7a) and (C7b) to arrive at Eqs. (3a) and (3b) presented in the main text.

## Appendix D: Generation and analysis of porous geometries

The simulation geometries have been created with *PoreSpy* [83], a freely available open-source python library. *PoreSpy* provides built-in functions for the generation of basic porous materials, such as the fibrous media used here, as well as diagnostic tools to, e.g., determine pore size and chord length distributions. The geometry outputs consist of binary voxel images (*TIFF* format) that partition the domain into void and fiber phases, making them readily integrable into the CPM framework. The porous geometries used in this work are available as *TIFF* files on [GitHub](https://github.com/GrawLab/CellMigration_Porosity).

## Footnotes

1 We apply the mean-field approximation ⟨ln *x*⟩ *≈* ln⟨*x*⟩.

## References

[1] U. H. von Andrian and T. R. Mempel, Homing and cellular traffic in lymph nodes, Nature Reviews. Immunology 3, 867 (2003).

[2] M. Zhao, B. Song, J. Pu, T. Wada, B. Reid, G. Tai, F. Wang, A. Guo, P. Walczysko, Y. Gu, T. Sasaki, A. Suzuki, J. V. Forrester, H. R. Bourne, P. N. Devreotes, C. D. McCaig, and J. M. Penninger, Electrical signals control wound healing through phosphatidylinositol-3-OH kinase-γ and PTEN, Nature 442, 457 (2006).

[3] D. Masopust and J. M. Schenkel, The integration of T cell migration, differentiation and function, Nature Reviews Immunology 13, 309 (2013).

[4] R. Ananthakrishnan and A. Ehrlicher, The Forces Behind Cell Movement, International Journal of Biological Sciences, 303 (2007).

[5] M. Murrell, P. W. Oakes, M. Lenz, and M. L. Gardel, Forcing cells into shape: The mechanics of actomyosin contractility, Nature Reviews Molecular Cell Biology 16, 486 (2015).

[6] W.-J. Rappel and L. Edelstein-Keshet, Mechanisms of cell polarization, Current Opinion in Systems Biology 3, 43 (2017).

[7] K. M. Yamada and M. Sixt, Mechanisms of 3D cell migration, Nature Reviews Molecular Cell Biology 20, 738 (2019).

[8] F. Merino-Casallo, M. J. Gomez-Benito, S. Hervas-Raluy, and J. M. Garcia-Aznar, Unravelling cell migration: Defining movement from the cell surface, Cell Adhesion & Migration 16, 25 (2022).

[9] M. R. Chastney, J. Kaivola, V.-M. Leppänen, and J. Ivaska, The role and regulation of integrins in cell migration and invasion, Nature Reviews Molecular Cell Biology 26, 147 (2025).

[10] X. Trepat, Z. Chen, and K. Jacobson, Cell Migration, Comprehensive Physiology 2, 2369 (2012).

[11] S. SenGupta, C. A. Parent, and J. E. Bear, The principles of directed cell migration, Nature Reviews Molecular Cell Biology 22, 529 (2021).

[12] J. d’Alessandro, A. Barbier–Chebbah, V. Cellerin, O. Benichou, R. M. Mège, R. Voituriez, and B. Ladoux, Cell migration guided by long-lived spatial memory, Nature Communications 12, 4118 (2021).

[13] J. Renkawitz, A. Kopf, J. Stopp, I. De Vries, M. K. Driscoll, J. Merrin, R. Hauschild, E. S. Welf, G. Danuser, R. Fiolka, and M. Sixt, Nuclear positioning facilitates amoeboid migration along the path of least resistance, Nature 568, 546 (2019).

[14] Z. Sadjadi, R. Zhao, M. Hoth, B. Qu, and H. Rieger, Migration of Cytotoxic T Lymphocytes in 3D Collagen Matrices, Biophysical Journal 119, 2141 (2020).

[15] T. H. Harris, E. J. Banigan, D. A. Christian, C. Konradt, E. D. Tait Wojno, K. Norose, E. H. Wilson, B. John, W. Weninger, A. D. Luster, A. J. Liu, and C. A. Hunter, Generalized Lévy walks and the role of chemokines in migration of effector CD8+ T cells, Nature 486, 545 (2012).

[16] T. R. Mempel, T. Junt, and U. H. Von Andrian, Rulers over Randomness: Stroma Cells Guide Lymphocyte Migration in Lymph Nodes, Immunity 25, 867 (2006).

[17] G. M. Fricke, K. A. Letendre, M. E. Moses, and J. L. Cannon, Persistence and Adaptation in Immunity: T Cells Balance the Extent and Thoroughness of Search, PLOS Computational Biology 12, e1004818 (2016).

[18] P.-H. Wu, A. Giri, S. X. Sun, and D. Wirtz, Three-dimensional cell migration does not follow a random walk, Proceedings of the National Academy of Sciences 111, 3949 (2014).

[19] J. W. Griffith, C. L. Sokol, and A. D. Luster, Chemokines and Chemokine Receptors: Positioning Cells for Host Defense and Immunity, Annual Review of Immunology 32, 659 (2014).

[20] O. Schulz, S. I. Hammerschmidt, G. L. Moschovakis, and R. Förster, Chemokines and Chemokine Receptors in Lymphoid Tissue Dynamics, Annual Review of Immunology 34, 203 (2016).

[21] K. J. Cheung and S. Horne-Badovinac, Collective migration modes in development, tissue repair and cancer, Nature Reviews Molecular Cell Biology 10.1038/s41580-025-00858-9 (2025).

[22] M. Miron-Mendoza, J. Seemann, and F. Grinnell, The differential regulation of cell motile activity through matrix stiffness and porosity in three dimensional collagen matrices, Biomaterials 31, 6425 (2010).

[23] S. R. Peyton, Z. I. Kalcioglu, J. C. Cohen, A. P. Runkle, K. J. Van Vliet, D. A. Lauffenburger, and L. G. Griffith, Marrow-Derived stem cell motility in 3D synthetic scaffold is governed by geometry along with adhesivity and stiffness, Biotechnology and Bioengineering 108, 1181 (2011).

[24] A. Pathak and S. Kumar, Independent regulation of tumor cell migration by matrix stiffness and confinement, Proceedings of the National Academy of Sciences 109, 10334 (2012).

[25] K. Wolf, M. Te Lindert, M. Krause, S. Alexander, J. Te Riet, A. L. Willis, R. M. Hoffman, C. G. Figdor, S. J. Weiss, and P. Friedl, Physical limits of cell migration: Control by ECM space and nuclear deformation and tuning by proteolysis and traction force, Journal of Cell Biology 201, 1069 (2013).

[26] J.-P. Bouchaud and A. Georges, Anomalous diffusion in disordered media: Statistical mechanisms, models and physical applications, Physics Reports 195, 127 (1990).

[27] V. Zaburdaev, S. Denisov, and J. Klafter, Lévy walks, Reviews of Modern Physics 87, 483 (2015).

[28] J. Bickmann and R. Wittkowski, Collective dynamics of active Brownian particles in three spatial dimensions: A predictive field theory, Physical Review Research 2, 033241 (2020).

[29] A. Datta, C. Beta, and R. Großmann, Random walks of intermittently self-propelled particles, Physical Review Research 6, 043281 (2024).

[30] G. S. Giardini, G. L. Thomas, C. R. Da Cunha, and R. M. De Almeida, Membrane fluctuations in migrating mesenchymal cells preclude instantaneous velocity definitions, Physica A: Statistical Mechanics and its Applications 647, 129915 (2024).

[31] J. B. Beltman, A. F. Marée, J. N. Lynch, M. J. Miller, and R. J. De Boer, Lymph node topology dictates T cell migration behavior, The Journal of Experimental Medicine 204, 771 (2007).

[32] M. Chiang and D. Marenduzzo, Glass transitions in the cellular Potts model, EPL (Europhysics Letters) 116, 28009 (2016).

[33] B. Loewe, M. Chiang, D. Marenduzzo, and M. C. Marchetti, Solid-Liquid Transition of Deformable and Overlapping Active Particles, Physical Review Letters 125, 038003 (2020).

[34] L. Van Steijn, J. A. Wondergem, K. Schakenraad, D. Heinrich, and R. M. Merks, Deformability and collision-induced reorientation enhance cell topotaxis in dense microenvironments, Biophysical Journal 122, 2791 (2023).

[35] A. Hopkins, B. Loewe, M. Chiang, D. Marenduzzo, and M. C. Marchetti, Motility induced phase separation of deformable cells, Soft Matter 19, 8172 (2023).

[36] D. B. Brückner, A. Fink, C. Schreiber, P. J. F. Röttgermann, J. O. Radler, and C. P. Broedersz, Stochastic nonlinear dynamics of confined cell migration in two-state systems, Nature Physics 15, 595 (2019).

[37] D. B. Brückner and C. P. Broedersz, Learning dynamical models of single and collective cell migration: A review, Reports on Progress in Physics 87, 056601 (2024).

[38] T. Brandstätter, E. Brieger, D. B. Brückner, G. Ladurner, J. O. Radler, and C. P. Broedersz, Data-Driven Theory Reveals Protrusion and Polarity Interactions Governing Collision Behavior of Distinct Motile Cells, PRX Life 3, 033015 (2025).

[39] K. Goswami, A. G. Cherstvy, A. Godec, and R. Metzler, Anomalous diffusion of active Brownian particles in responsive elastic gels: Nonergodicity, non-Gaussianity, and distributions of trapping times, Physical Review E 110, 044609 (2024).

[40] K. Schakenraad, L. Ravazzano, N. Sarkar, J. A. J. Wondergem, R. M. H. Merks, and L. Giomi, Topotaxis of active Brownian particles, Physical Review E 101, 032602 (2020).

[41] F. Moore, J. Russo, T. B. Liverpool, and C. P. Royall, Active Brownian particles in random and porous environments, The Journal of Chemical Physics 158, 104907 (2023).

[42] M. S. Alber, M. A. Kiskowski, J. A. Glazier, and Y. Jiang, On Cellular Automaton Approaches to Modeling Biological Cells, in Mathematical Systems Theory in Biology, Communications, Computation, and Finance, Vol. 134, edited by D. N. Arnold, F. Santosa, J. Rosenthal, and D. S. Gilliam (Springer New York, 2003) pp. 1–39.

[43] T. Hirashima, E. G. Rens, and R. M. H. Merks, Cellular Potts modeling of complex multicellular behaviors in tissue morphogenesis, Development, Growth & Differentiation 59, 329 (2017).

[44] F. Ziebert and I. S. Aranson, Computational approaches to substrate-based cell motility, npj Computational Materials 2, 16019 (2016).

[45] A. Moure and H. Gomez, Phase-field model of cellular migration: Three-dimensional simulations in fibrous networks, Computer Methods in Applied Mechanics and Engineering 320, 162 (2017).

[46] A. Moure and H. Gomez, Phase-Field Modeling of Individual and Collective Cell Migration, Archives of Computational Methods in Engineering 28, 311 (2021).

[47] M. E. Cates and J. Tailleur, Motility-Induced Phase Separation, Annual Review of Condensed Matter Physics 6, 219 (2015).

[48] D. Bi, X. Yang, M. C. Marchetti, and M. L. Manning, Motility-Driven Glass and Jamming Transitions in Biological Tissues, Physical Review X 6, 021011 (2016).

[49] M. R. Shaebani, A. Wysocki, R. G. Winkler, G. Gompper, and H. Rieger, Computational models for active matter, Nature Reviews Physics 2, 181 (2020).

[50] A. Ziepke, I. Maryshev, I. S. Aranson, and E. Frey, Multiscale organization in communicating active matter, Nature Communications 13, 6727 (2022).

[51] A. Hayn, T. Fischer, and C. T. Mierke, Inhomogeneities in 3D Collagen Matrices Impact Matrix Mechanics and Cancer Cell Migration, Frontiers in Cell and Developmental Biology 8, 593879 (2020).

[52] C. Kurzthaler, S. Mandal, T. Bhattacharjee, H. Löwen, S. S. Datta, and H. A. Stone, A geometric criterion for the optimal spreading of active polymers in porous media, Nature Communications 12, 7088 (2021).

[53] S. Torquato and B. Lu, Chord-length distribution function for two-phase random media, Physical Review E 47, 2950 (1993).

[54] T. Bhattacharjee and S. S. Datta, Bacterial hopping and trapping in porous media, Nature Communications 10, 2075 (2019).

[55] V. N. Burganos and S. V. Sotirchos, Simulation of Knudsen diffusion in random networks of parallel pores, Chemical Engineering Science 43, 1685 (1988).

[56] M. M. Tomadakis and S. V. Sotirchos, Effective Kundsen diffusivities in structures of randomly overlapping fibers, AIChE Journal 37, 74 (1991).

[57] P. L. Krapivsky, S. Redner, and E. Ben-Naim, A Kinetic View of Statistical Physics (Cambridge University Press, 2010).

[58] J. Han, S. W. Turner, and H. G. Craighead, Entropic Trapping and Escape of Long DNA Molecules at Submicron Size Constriction, Physical Review Letters 83, 1688 (1999).

[59] R. Zwanzig, Diffusion past an entropy barrier, The Journal of Physical Chemistry 96, 3926 (1992).

[60] A.-L. Barabási and M. Pósfai, Network Science (Cambridge University Press, 2016).

[61] H. Kramers, Brownian motion in a field of force and the diffusion model of chemical reactions, Physica 7, 284 (1940).

[62] S. N. Mueller, M. Matloubian, D. M. Clemens, A. H. Sharpe, G. J. Freeman, S. Gangappa, C. P. Larsen, and R. Ahmed, Viral targeting of fibroblastic reticular cells contributes to immunosuppression and persistence during chronic infection, Proceedings of the National Academy of Sciences 104, 15430 (2007).

[63] J. L. Chitty and T. R. Cox, The extracellular matrix in cancer: From understanding to targeting, Trends in Cancer, S2405803325001268 (2025).

[64] A. Pacheco-Pozo, M. Balcerek, A. Wy-lomanska, K. Burnecki, I. M. Sokolov, and D. Krapf, Langevin Equation in Heterogeneous Landscapes: How to Choose the Interpretation, Physical Review Letters 133, 10.1103/phys-revlett.133.067102 (2024).

[65] T. Bhattacharjee and S. S. Datta, Confinement and activity regulate bacterial motion in porous media, Soft Matter 15, 9920 (2019).

[66] A. Datta, S. Beier, V. Pfeifer, R. Großmann, and C. Beta, Bacterial swimming in porous gels exhibits intermittent run motility with active turns and mechanical trapping, Scientific Reports 15, 20320 (2025).

[67] C. Bousige, P. Levitz, and B. Coasne, Bridging scales in disordered porous media by mapping molecular dynamics onto intermittent Brownian motion, Nature Communications 12, 1043 (2021).

[68] T. Bhattacharjee, D. B. Amchin, J. A. Ott, F. Kratz, and S. S. Datta, Chemotactic migration of bacteria in porous media, Biophysical Journal 120, 3483 (2021).

[69] R. Sunyer and X. Trepat, Durotaxis, Current Biology 30, R383 (2020).

[70] J. Park, D.-H. Kim, H.-N. Kim, C. J. Wang, M. K. Kwak, E. Hur, K.-Y. Suh, S. S. An, and A. Levchenko, Directed migration of cancer cells guided by the graded texture of the underlying matrix, Nature Materials 15, 792 (2016).

[71] R. K. Sadhu, M. Luciano, W. Xi, C. Martinez-Torres, M. Schröder, C. Blum, M. Tarantola, S. Villa, S. Penič, Iglič, C. Beta, O. Steinbock, E. Bodenschatz, Ladoux, S. Gabriele, and N. S. Gov, A minimal physical model for curvotaxis driven by curved protein complexes at the cell’s leading edge, Proceedings of the National Academy of Sciences 121, e2306818121 (2024).

[72] D. Fuller, W. Chen, M. Adler, A. Groisman, H. Levine, W.-J. Rappel, and W. F. Loomis, External and internal constraints on eukaryotic chemotaxis, Proceedings of the National Academy of Sciences 107, 9656 (2010).

[73] G. Micali and R. G. Endres, Bacterial chemotaxis: Information processing, thermodynamics, and behavior, Current Opinion in Microbiology 30, 8 (2016).

[74] I. Andreu, B. Falcones, S. Hurst, N. Chahare, X. Quiroga, A.-L. Le Roux, Z. Kechagia, A. E. M. Beedle, A. Elosegui-Artola, X. Trepat, R. Farré, T. Betz, I. Almendros, and P. Roca-Cusachs, The force loading rate drives cell mechanosensing through both reinforcement and cytoskeletal softening, Nature Communications 12, 4229 (2021).

[75] A. Pathni, K. Wagh, I. Rey-Suarez, and A. Upadhyaya, Mechanical regulation of lymphocyte activation and function, Journal of Cell Science 137, jcs219030 (2024).

[76] S. S. Deville and N. Cordes, The Extracellular, Cellular, and Nuclear Stiffness, a Trinity in the Cancer Resistome—A Review, Frontiers in Oncology 9, 1376 (2019).

[77] R. Borst, L. Meyaard, and M. I. Pascoal Ramos, Understanding the matrix: Collagen modifications in tumors and their implications for immunotherapy, Journal of Translational Medicine 22, 382 (2024).

[78] M. F. Krummel, F. Bartumeus, and A. Gérard, T cell migration, search strategies and mechanisms, Nature Reviews Immunology 16, 193 (2016).

[79] H. Du, J. M. Bartleson, S. Butenko, V. Alonso, W. F. Liu, D. A. Winer, and M. J. Butte, Tuning immunity through tissue mechanotransduction, Nature Reviews Immunology 23, 174 (2023).

[80] B. D. Hale, Y. Severin, F. Graebnitz, D. Stark, D. Guignard, J. Mena, Y. Festl, S. Lee, J. Hanimann, N. S. Zangger, M. Meier, D. Goslings, O. Lamprecht, B. M. Frey, A. Oxenius, and B. Snijder, Cellular architecture shapes the näive T cell response, Science 384, eadh8697 (2024).

[81] J. Starruß, W. De Back, L. Brusch, and A. Deutsch, Morpheus: A user-friendly modeling environment for multiscale and multicellular systems biology, Bioinformatics 30, 1331 (2014).

[82] C. W. Gardiner, Handbook of Stochastic Methods for Physics, Chemistry, and the Natural Sciences, Springer Series in Synergetics No. v. 13 (Springer-Verlag, 1983).

[83] J. Gostick, Z. Khan, T. Tranter, M. Kok, M. Agnaou, M. Sadeghi, and R. Jervis, PoreSpy: A Python Toolkit for Quantitative Analysis of Porous Media Images, Journal of Open Source Software 4, 1296 (2019).

